# Bent DNA bows as amplifiers and biosensors for detecting DNA-interacting salts and molecules

**DOI:** 10.1101/2020.02.19.956144

**Authors:** Jack Freeland, Lihua Zhang, Shih-Ting Wang, Mason Ruiz, Yong Wang

## Abstract

Due to the central role of DNA, its interactions with inorganic salts and small organic molecules are important for understanding various fundamental cellular processes in living systems, deciphering the mechanism of many diseases related to DNA damages, and discovering or designing inhibitors and drugs targeting DNA. However, there is still a need for improved sensitivity to detect these interactions, especially in situations where expensive sophisticated equipment is not available. Here we report our development and demonstration of bent DNA bows for amplifying, sensing, and detecting the interactions of 14 inorganic salts and small organic molecules with DNA. With the bent DNA bows, these interactions were easily visualized and quantified in gel electrophoresis, which were difficult to measure without bending. In addition, the strength of the interactions of DNA with the various salts/molecules were quantified using the modified Hill equation. This work highlights the amplification effects of the bending elastic energy stored in the DNA bows and the potential use of the DNA bows for quantitatively measuring DNA interactions with small molecules as simple economic methods; it may also pave the way for exploiting the bent DNA bows for other applications such as monitoring water quality and screening DNA-targeting molecules and drugs.

## Introduction

DNA is one of the most essential elements of life, and its interactions with inorganic salts and small organic molecules are important for many reasons ^1,2^. First, understanding these interactions is critical for understanding various fundamental cellular processes in living systems ^1,3–5^. For example, genomic stability and DNA repair rely on the presence, mediation, and/or participation of metal ions ^1,6–8^. Second, the DNA interactions with salts/molecules are important for the mechanism of diseases, especially those related to DNA damaging and repairing ^9–11^. For example, heavy metal ions and various chemical carcinogens and mutagens interact and react with DNA directly or indirectly, causing many human cancers ^12–14^. Third, understanding these DNA interactions helps to discover, design and develop inhibitors and drugs targeting DNA for treating various diseases ^15–17^. For example, DNA damaging agents have been widely used in treating many cancers ^16–18^. Therefore, it is important to understand the interactions between DNA and inorganic salts or small organic molecules.

Various techniques have been developed for investigating the interactions between DNA and other molecules, including gel electrophoresis, melting-curve, fluorescence anisotropy, circular dichroism, isothermal titration calorimetry, x-ray absorption spectroscopy, optical tweezers, magnetic tweezers, electron paramagnetic resonance, Raman spectroscopy, and nuclear magnetic resonance spectroscopy ^19–37^. However, there is still a need for improved sensitivity to detect these interactions. First, improving the sensitivity helps to identify molecules that interact with DNA weakly ^38^. Second, when expensive sophisticated equipment is not available, there is a need to develop a general strategy to enhance the sensitivities of detection of weak interactions of DNA with various molecules ^38^.

In this work, we demonstrated the application of bent DNA bows ^39–42^ as amplifiers and sensors for detecting and measuring the interactions between DNA and 14 different inorganic salts and small organic molecules. The DNA bows are constructed as illustrated in Fig. 1A and described in the “Material and Methods” section. They consist of two segments: a bent double-stranded segment and a stretched single-stranded part ^39–42^. The mechanical energy stored in the bent DNA bows makes them more susceptible and sensitive to perturbations caused by the interactions of DNA with other molecules ^41,43,44^. For example, the bending elastic energy in the DNA bows helps to drive the formation of dimers (Fig. 1B) or higher-order oligomers or to stimulate the dissociation of double strands into single strands, as the elastic energy gets released in both the dimers or single strands ^39–42^. Due to the improved susceptibility and sensitivity, the bent DNA bows enabled us to detect these interactions much more easily than unbent DNA. In addition, we showed that this technique based on bent DNA bows were capable of quantifying these interactions by fitting the relationship between the percentage of bent DNA bows and the concentrations of the tested salts or molecules presented in the solutions using the modified Hill equations, which report the strength of the DNA-salt or DNA-organic compound interactions.

**Figure 1.**
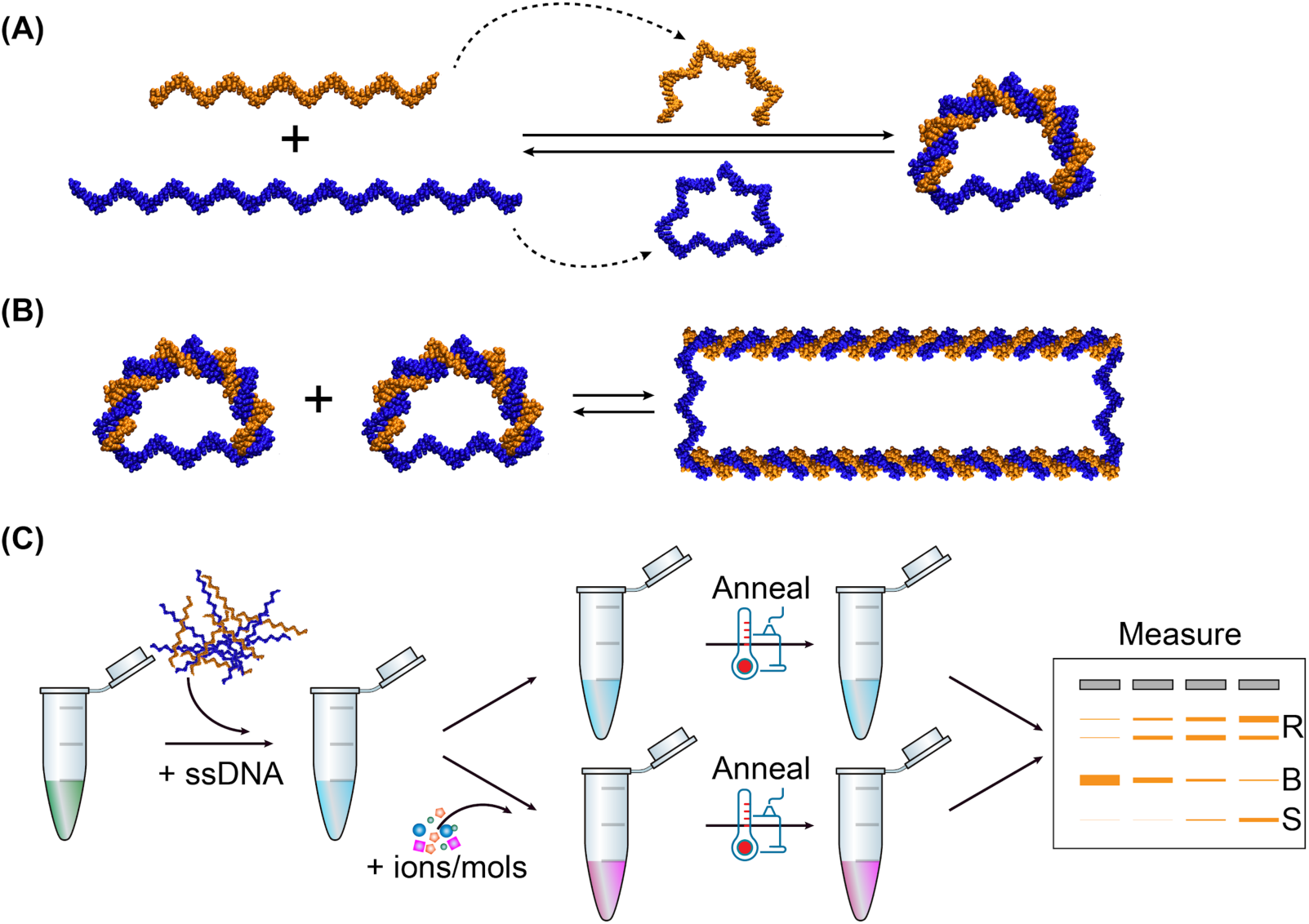
Illustration of biosensors based on bent DNA bows. (A) Construction of DNA bows from synthesized single-stranded DNA. (B) Relaxation of bending elastic energy in DNA bows by forming dimers. (C) Sketch of the procedure for detecting DNA interactions with ions and molecules, visualized by gel electrophoresis as an example.

## Material and Methods

### Construction of DNA bows

Synthetic DNA oligos were purchased from Integrated DNA Technologies (IL, USA) and resuspended in distilled water to a final concentration of 100 μM. The oligos used in this study include S45 (5’ - CAC AGA ATT CAG CAG CAG GCA ATG ACA GTA GAC ATA CGA CGA CTC -3’), S30B (5’ - CTG CTG AAT TCT GTG GAG TCG TCG TAT GTC - 3’), S45CF (5’ - GAG TCG TCG TAT GTC TAC TGT CAT TGC CTG CTG CTG AAT TCT GTG - 3’), S30CM (5’ - GTA TGT CTA CTG TCA TTG CCT GCT GCT GAA - 3’), and S30CR (5’ - TAC TGT CAT TGC CTG CTG CTG AAT TCT GTG - 3’).

Bent DNA bows were constructed from two synthesized single-stranded DNA (S45 and S30B) via self-assembly following our previous work ^41^. The sequences of the single strands were designed such that the last 15 bases at the 5’-end of the long strand (S45) hybridize to the 5’-half of the short strand (S30B), while the last 15 bases at the 3’-end of S45 hybridize to the 3’-half of S30B. Upon hybridization, a circular construct is formed, with a double-stranded segment of 30 base pairs (bp) with a nick and a single-stranded segment of 15 bases ^39–42^. Three linear constructs (CF, CM, and CR, shown in Fig. S1) were used as negative controls. Upon hybridization, CF (S45 + S45CF) is double-stranded completely, while CM (S45 + S30CM) and CR (S45 + S30CR) have overhangs of single strands at one or two sides, respectively. The long strands for CF and CM are the same as the long one in the DNA bows.

### Detection of DNA-interacting salts/molecules using DNA bows

The detection of DNA-interacting salts/molecules using DNA bows is illustrated in Fig. 1C. Briefly, single strands (S45 and S30B) were mixed at equal molar amounts in background buffer (0.4 mM Tris-HCl with pH adjusted to 7.5 and 0.5 mM NaCl) to reach a final concentration of 2 μM ^41^. The DNA samples without other ions or molecules of interest were used as baselines/controls. To detect possible interactions of ions and molecules of interest, solutions of the salts/molecules of interest were prepared in water and mixed with DNA strands in the background buffer to reach desired concentrations (Table 1). The mixtures were heated to 75ºC for 2 min and gradually cooled down to 22ºC (room temperature) in 5 hr ^41^. The mixtures were incubated at 22ºC for overnight to allow full equilibrium, followed by gel electrophoresis for visualization on the second day ^41^.

**Table 1.** Salts / molecules and their concentrations used in this study.

| Salt / Molecule | Concentrations |
| --- | --- |
| MgCl<sub>2</sub> | 0, 1, 2, 3, 4, 5, 6, 7 mM |
| MgSO<sub>4</sub> | 0, 1, 2, 3, 4, 5, 6, 7 mM |
| KCl | 0, 1, 2, 3, 4, 5, 6, 7 mM |
| CaCl<sub>2</sub> | 0, 1, 2, 3, 4, 5, 6, 7 mM |
| Al(NO<sub>3</sub>)<sub>3</sub> | 0, 16, 32, 48, 64, 80, 96, 112 µM |
| Zn(NO<sub>3</sub>)<sub>2</sub> | 0, 30, 60, 90, 120, 150, 180, 210 µM |
| AgNO<sub>3</sub> | 0, 10, 20, 30, 40, 50, 60, 70, 80, 90 µM |
| Guanidine | 0, 1, 2, 3, 4, 5, 6, 7 mM |
| Putrescine | 0, 0.5, 1, 2, 4, 8, 16, 32 mM |
| Spermidine | 0, 6, 12, 18, 24, 30, 36, 42 µM |
| Ganciclovir | 0, 1, 2, 3, 4, 5, 6, 7 mM |
| Thiamine | 0, 36, 72, 108, 144, 180, 300, 600 µM |
| Ethidium Bromide | 0, 381, 400, 419, 438, 457, 476, 495 µM |
| SYBR Safe * | 0, 25, 50, 60, 70, 80, 90, 100 µM |

\* It is assumed that the concentration of 1X SYBR Safe (commercially available from Thermo Fisher Scientific) is 1 µM, according to the corresponding patent <sup>45</sup>.

### Gel electrophoresis

Polyacrylamide gels (12%) were prepared in the laboratory. Briefly, 3 mL of acrylamide/bis solutions (40%, Bio-Rad Laboratories, CA, USA), 1 mL of 10X tris-borate-EDTA (TBE) buffer (Bio-Rad Laboratories), 20 μL of freshly made ammonium persulfate (APS, 10% in water, Thermo Fisher Scientific, MA, USA) and 6 mL of distilled water were mixed thoroughly and degassed for 5 min in vacuum. The mixture was poured into gel cast cassette immediately after adding 10 μL of tetramethylethylenediamine (TEMED) (Thermo Fisher Scientific), followed by incubation at room temperature for 20 – 60 min to allow full gelation before use.

The prepared DNA samples (5 μL) were mixed thoroughly with 1 μL of 6X DNA loading buffer (Bio-Rad Laboratories). The mixtures were loaded into the wells of the prepared gel. The gel electrophoresis (Edvotek Inc., DC, USA) was run at 100V for 50 – 60 min in 1X TBE buffer, followed by staining the gel with 1X SYBR Safe solution (Thermo Fisher Scientific) for 15 – 30 min with gentle shaking. The stained gel was then imaged with a typical exposure time of 1 – 5 sec using a gel documentation system (Analytik Jena US LLC, CA, USA).

The acquired gel images were analyzed using ImageJ ^46,47^. The original gel images were first rotated, cropped, and inverted, followed by subtracting the background with a rolling ball radius of 10 pixels ^48^. The preprocessed gel lanes were then analyzed using the gel analysis procedure in ImageJ ^46,47^, from which the intensities of the bands of DNA bows and that of the bands corresponding to the relaxed ones (dimers, trimers, and oligomers) were obtained. Lastly, the intensities were rescaled by dividing the band intensity of the DNA bows on the same gel in the absence of ions/molecules of interest ^41^.

### Visualization of DNA bows using transmission electron microscopy (TEM)

The morphology of the bent DNA bows was visualized by TEM imaging. Briefly, the prepared DNA bows (∼2 μM) were diluted in the background buffer to a final concentration of ∼200 nM. Then, 5 μL of the diluted DNA solution was dropped onto a carbon film coated TEM grid (Electron Microscopy Sciences, PA, USA), and incubated at room temperature for 1.5 min, followed by removing the residual liquid with filter papers. The grid was washed by 5 μL of deionized water and stained with 5 μL of 2 wt% Nano-W^TM^ (Nanoprobes Inc., NY, USA) for 10 sec ^49^. After removing excess liquid with filter papers, the DNA sample was imaged using a JEOL 2100F TEM with an acceleration voltage of 200 kV.

## Results and Discussions

### Visualization of DNA bows using TEM

TEM imaging was performed to directly visualize the constructed DNA bows after negative staining with organo-tungstate compounds (Nano-W^TM^) ^49^. Examples of TEM images of individual DNA bows clearly showed the bending structures (Fig. 2A), which are presumably the bent, double-stranded part of the DNA bows. For confirmation, the arc lengths of the bending structures on the TEM images were quantified using ImageJ ^46,47^. A single peak was observed in the distribution of the measured arc lengths (Fig. 2B). Fitting the peak with the Gaussian distribution resulted in an average length of 8.4 ± 1.4 nm (mean ± standard deviation). Considering that the double-stranded segment of DNA bows has a length of 30 bp, the measured arc length of the DNA bows was consistent with previous reports from direct TEM imaging of DNA structures and X-ray data ^50^, confirming that the dark bending structures in the TEM images were indeed bent, double-stranded part of the DNA bows.

**Figure 2.**
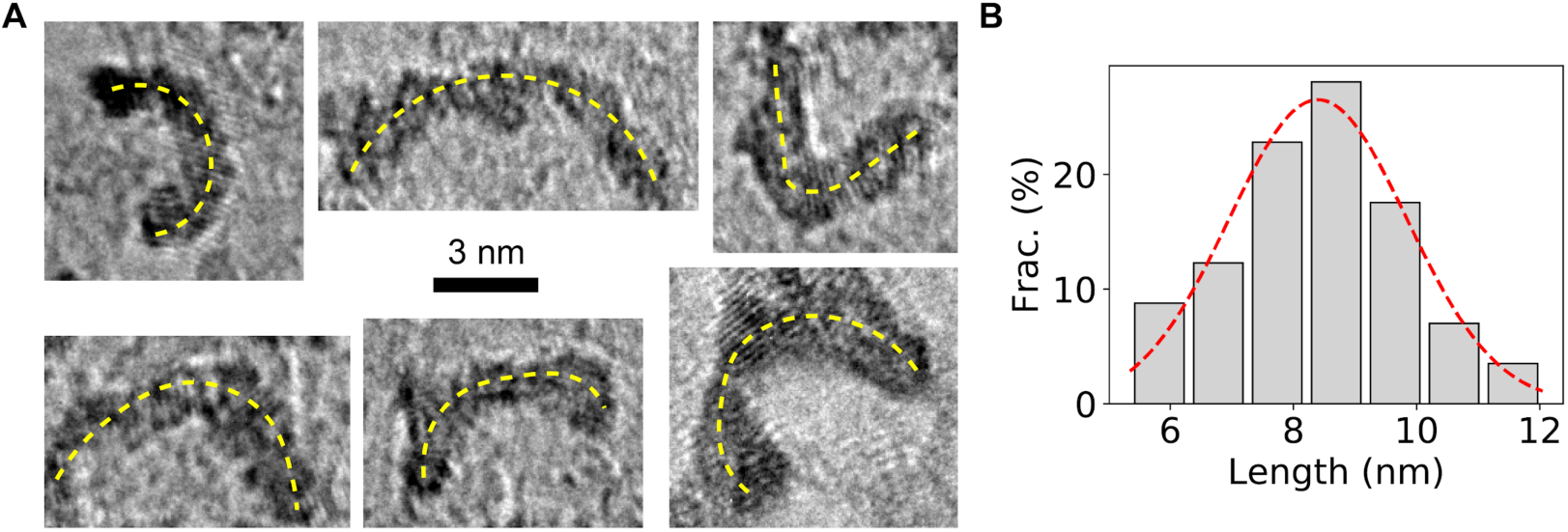
TEM visualization of bent DNA bows. (A) Examples of TEM images of DNA bows. The bending of the DNA bows is highlighted by the yellow dashed lines. Scale bar = 3 nm. (B) Distribution of the arc lengths of DNA bows, which was fitted by a Gaussian distribution (red dashed line), resulting in an average length of 8.4 ± 1.4nm (mean ± standard deviation).

### Detection of inorganic ions using DNA bows

The DNA bows were applied to detect the interactions of DNA with various inorganic ions. It is well-known that these DNA-ion interactions play essential roles in the properties and functions of DNA molecules ^1,8^. One of the most famous examples is the local and long-range electrostatic interactions of cations on the structure and stiffness of DNA ^19,21,51^. In particular, Mg^2+^ ions are well-known to mediate and stabilize the secondary structures of DNA, playing critical roles in genomic packaging, gene regulation, and DNA-repairing ^19,21,51,52^. On the other hand, certain ions, especially heavy metal ions such as Al^3+^, Ag^+^, and Zn^2+^, can damage DNA molecules, accumulation of which is associated with various diseases including cancers ^15–17^.

We first examined the interaction of DNA with Mg^2+^ ions from two different salts, MgCl_2_ and MgSO_4_, and found that both salts were capable of driving the formation of relaxed DNA loops (Fig. 3A and 3B). In the absence of the bent DNA bows to amplify the signals (i.e., with linear DNA controls as shown in Fig. S1), the DNA molecules treated with MgCl_2_ and MgSO_4_ salts at concentrations up to 7 mM did not show any observable difference in gel electrophoresis (Fig. S2, rows indicated by “CF”, “CM”, and “CR”). Quantifying the intensities of the bands showed little differences for the increasing concentrations of Mg^2+^ ions (◁, ▷, and 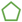 in Fig. 3A and 3B). Interestingly, when amplifying the signal of the DNA interactions with the MgCl_2_ and MgSO_4_ salts using the bent DNA bows, changes in the gel electrophoretic patterns of the DNA molecules were clear (Fig. 3A and 3B, and rows indicated by “Bent” in Fig. S2). The intensities of the DNA bows decreased as the concentrations of the Mg^2+^ salts increased (• in Fig. 3A and 3B). In addition, heavier bands corresponding to relaxed DNA loops (such as dimers, trimers, and/or oligomers) appeared in the presence of Mg^2+^ salts (insets of Fig. 3A and 3B) ^41^. By quantifying the intensities of the relaxed species (i.e., all other bands except the bands of DNA bows), we confirmed that the intensity of relaxed DNA loops increased, reaching a plateau at Mg^2+^ concentration ≈ 3 mM (□ in Fig. 3A and 3B).

**Figure 3.**
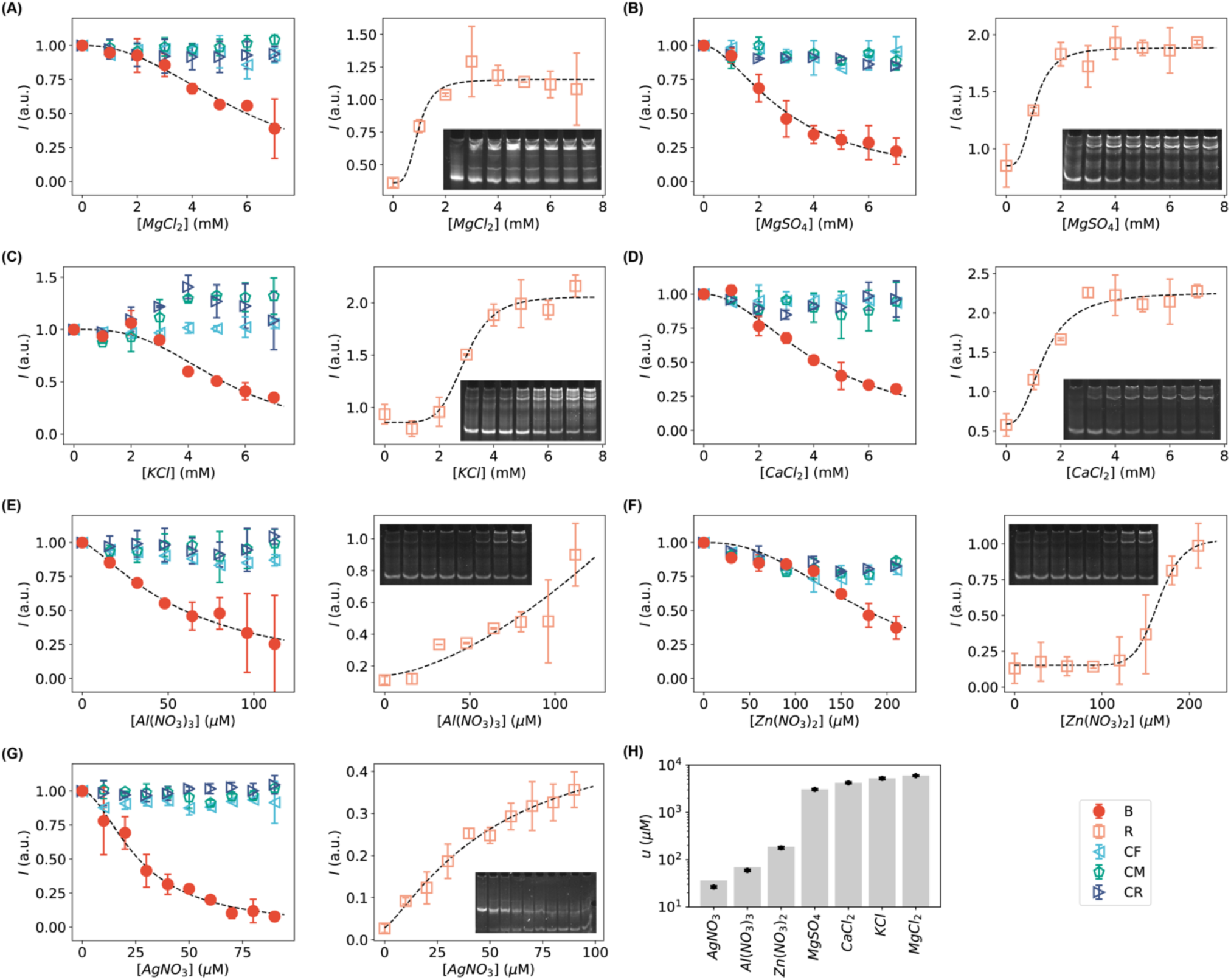
Intensities of the bands of DNA bows (red circles), relaxed DNA loops (orange squares), and linear DNA controls (cyan triangles, green pentagons, blue triangles) in the presence of various salts at increasing concentrations: (A) MgCl_2_, (B) MgSO_4_, (C) KCl, (D) CaCl_2_, (E) Al(NO_3_)_3_, (F) Zn(NO_3_)_2_ and (G) AgNO_3_. Insets are the representative, cropped gels of bent DNA bows in the presence of the corresponding salts at increasing concentrations. The corresponding full-length gels are shown in Fig. S2. (H) Fitted *u*-values for quantifying the strength of DNA interactions with inorganic salts.

In addition to the Mg^2+^-salts, we tested K^+^-salt (KCl) and Ca^2+^-salt (CaCl_2_) that benefits various cellular processes ^53–57^, and three other ions (Al^3+^, Zn^2+^, and Ag^+^, provided from the corresponding nitrate salts) that have been shown to be closely related to DNA damage *in vivo* ^15–17,58–62^. The K^+^ and Ca^2+^ ions resulted in similar effects on the DNA molecules in the same range of salt concentrations from 0 to 7 mM (Fig. 3 and 3D, and Fig. S2): the intensity of the band of the DNA bows decreased while the intensities of the relaxed heavier species increased. For the DNA-damaging ions, we observed that both Al^3+^ and Zn^2+^ ions resulted in the formation of heavier relaxed DNA loops, similar to Mg^2+^ ions; however, Ag^+^ ions led to dissociation of the bent DNA bows, as the band corresponding to the single-stranded DNA appeared ^41^. The difference in the change of gel patterns caused by the ions between Ag^+^ and all the other tested ions suggests that their interactions with DNA are distinct. Also note that the working concentrations of Al^3+^, Zn^2+^ and Ag^+^ ions were 10 – 100 times lower than that of Mg^2+^, K^+^, and Ca^2+^ ions.

To quantify the strength of the DNA-salt interactions by the bent DNA bows, we fitted the normalized intensities of the bands of DNA bows *I*_*B*_ as functions of the concentrations of the salts using an equation derived from the Hill equation that has been extensively used for characterizing the binding between ligands and macromolecules ^63,64^

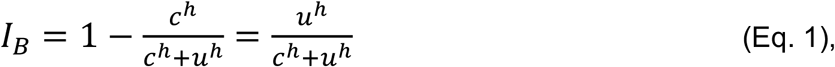

where *c* is the concentration of the tested salts, *h* is the Hill coefficient, and *u* is the characteristic concentration of the tested salts producing half intensity of the band of DNA bows in the absence of the salts. It turns out that (Eq. 1) fitted all the data very well (Fig. 3). The fitted parameters (*h* and *u*) were presented in Fig. S4A for the seven tested inorganic salts. In addition, the characteristic concentrations (*u*) for different salts were compared in Fig. 3H, which suggested that Al(NO_3_)_3_, Zn(NO_3_)_2_ and AgNO_3_ showed stronger interactions on DNA (i.e., lower *u* values) compared to MgCl_2_, MgSO_4_, KCl, and CaCl_2_.

In addition to the bands of DNA bows, we quantified the summed intensities of the relaxed bands *I* _R_and fitted them using the Hill equation ^63,64^ with the addition of a baseline (*b*),

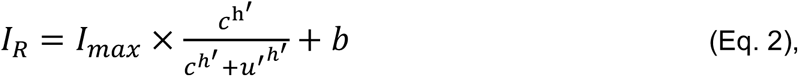

where *I*_max_ is the maximum intensity of the relaxed bands measured from the baseline. As shown in Fig. 3, the data from the relaxed bands can also be fitted well with the modified Hill equation, providing an additional way to quantify the interactions of the inorganic salts with DNA. Note that the two parameters in the Hill equation (*u*′ and *h*′) obtained from the relaxed bands are not necessarily the same as those estimated from the bands of DNA bows (*I*_B_), because the relaxed bands contain multiple relaxed species of different orders (i.e., dimers, trimers, tetramers, and other oligomers). Nonetheless, we found that *u*′ correlated very well with *u* for all the test inorganic salts, except for Al(NO_3_)_3_ (Fig. S5).

### Detection of small organic molecules using DNA bows

In addition to inorganic salts, we applied the bent DNA bows to amplify and detect the interactions of DNA with small organic molecules (Table 1). First, guanidine (guanidinium chloride, or guanidine hydrochloride, GuHCl) were tested as it is is a commonly used chaotropic agent at high concentrations to denature double-stranded DNA ^65,66^. Second, salts of putrescine and spermidine were chosen for testing our DNA bows because they have been reported previously to interact with DNA, decreasing the persistence length of DNA and making DNA softer ^21,67,68^. Third, we tested EtBr and SYBR safe, which are commonly used DNA intercalators and DNA staining dyes in gel electrophoresis ^69–71^. Lastly, we tested two molecules that are relevant to the production of DNA but unknown direct interactions with DNA: ganciclovir and thiamine. Ganciclovir is an antiviral medication used to treat cytomegalovirus (CMV) infections, and ganciclovir triphosphate is a competitive inhibitor of deoxyguanosine triphosphate (dGTP) incorporation into DNA ^72,73^. Thiamine is a vitamin that serves as a cofactor for a series of enzymes in different metabolic pathways and is required for the production of ATP, ribose, NAD, and DNA ^74,75^.

As expected, the interactions between DNA and guanidine, putrescine, spermidine, EtBr, or SYBR Safe could be amplified and detected by the bent DNA bows. It is noted that these interactions were not detectable or not significant without the bent DNA bow for amplification (i.e., with linear double-stranded DNA controls) at low enough concentrations of these organic molecules (rows indicated by “CF”, “CM”, and “CR” in Fig. S3). In contrast, when amplifying the signal of the DNA interactions with these organic molecules using the bent DNA molecules, the effects of the molecules at the same concentrations were observed (Fig. 4A-4E, and rows indicated by “Bent” in Fig. S3). Similar to the inorganic salts, the intensities of the bent DNA bands decreased as the concentrations of the organic molecules increased; however, the ranges of the “working” concentrations of the organic molecules were more diverse than the inorganic salts. It is noted that this diversity suggests that the observed interactions of organic molecules with the DNA bows were unlikely due to the residue ions in the solutions of organic molecules. In addition, the appearance of the heavier bands suggested that guanidine, putrescine, spermidine, EtBr, and SYBR Safe caused the formation of relaxed dimers or oligomers (Fig. 4A-4E). It is worthwhile to point out that the patterns of the heavier bands are different for different organic molecules, and some of the patterns are distinct from that of the inorganic salts (e.g., Fig. 4C-4E), which again suggests that the interactions of DNA with different organic molecules and inorganic salts are different.

**Figure 4.**
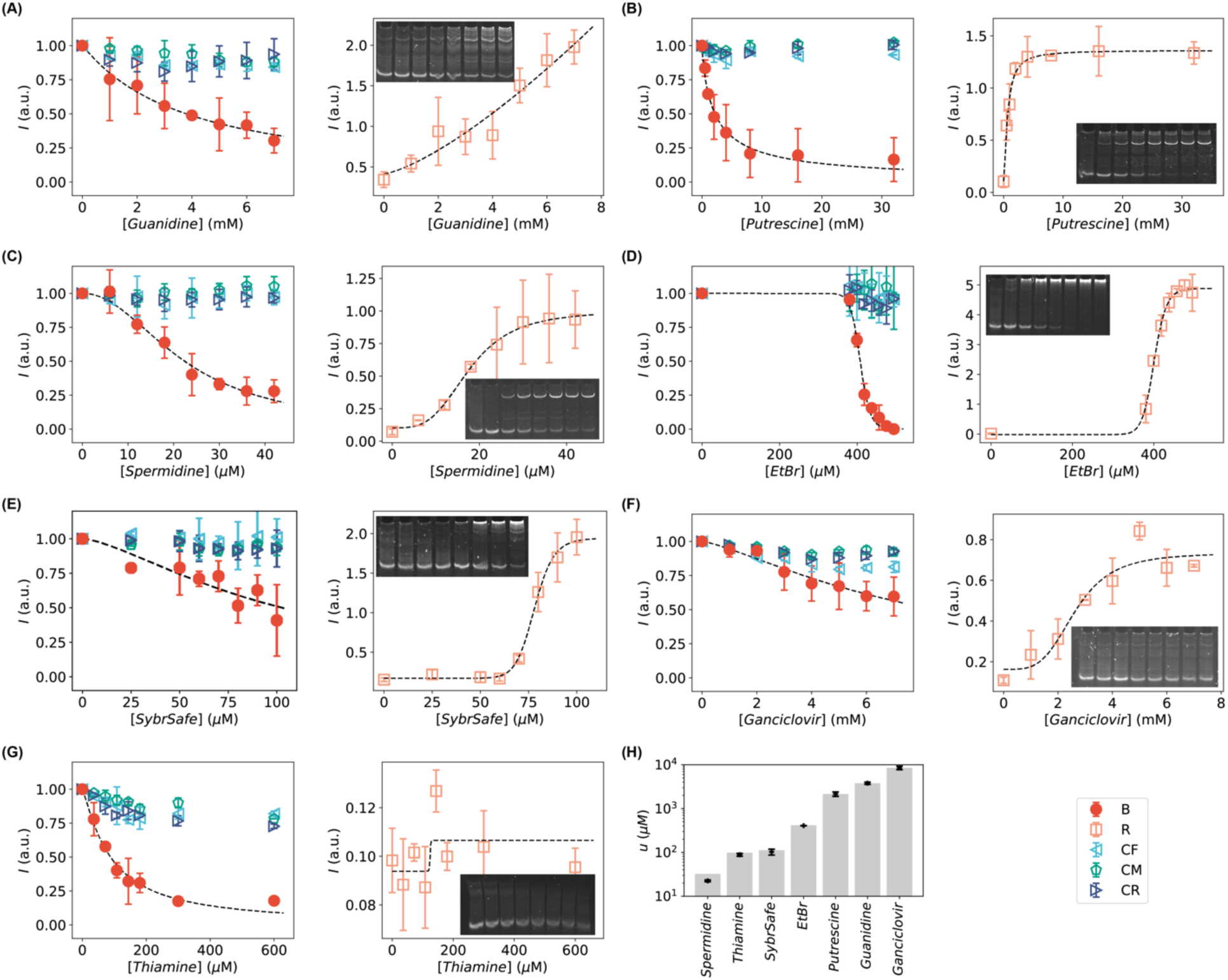
Intensities of the bands of DNA bows (red circles), relaxed DNA loops (orange squares), and linear DNA controls (cyan triangles, green pentagons, blue triangles) in the presence of small organic molecules at increasing concentrations: (A) guanidine, (B) putrescine, (C) spermidine, (D) ethidium bromide, (E) SYBR safe, (F) ganciclovir, and (G) thiamine. Insets are the representative, cropped gels of bent DNA bows in the presence of the corresponding small organic molecules at increasing concentrations. The corresponding full-length gels are shown in Fig. S3. (H) Fitted *u*-values for quantifying the strength of DNA interactions with organic molecules.

Interestingly, we observed that our bent DNA bows were also able to amplify and detect the interactions of DNA with ganciclovir and thiamine. For ganciclovir, the intensities of the bent DNA bands decreased, while faint bands of heavier species emerged as the concentration of ganciclovir increased (Fig. 4F). In contrast, heavier bands were absent as the concentration of thiamine increased, even if the intensities of DNA bows decreased (Fig. 4G). The decrease cannot be attributed to the interference of thiamine with DNA staining using SYBR safe after running the gel, as the decrease in the linear double-stranded DNA controls was much weaker (Fig. 4G, and Fig. S3).

Similar to the inorganic salts, the strength of the DNA interactions with these organic molecules were quantified using the modified Hill equations (Eq. 1 for *I*_*B*_, and Eq. 2 for *I*_*R*_), which again fitted all the data very well (except *I*_*R*_ for thiamine), as shown in Fig. 4. The fitted parameters from *I*_B_ (*h* and *u*) were presented in Fig. S4B for the seven tested organic molecules. In addition, the characteristic concentrations (*u*) for different organic molecules were compared in Fig. 4H. Furthermore, a good correlation between *u*′ obtained from *I* and Eq. 2 correlated and *u* was observed for all the organic molecules (except guanidine), although the fitting error of *u*′ for thiamine is large (Fig. S5).

## Conclusions

In conclusion, we demonstrated the application of bent DNA bows as amplifiers and sensors for detecting and quantifying the interactions between DNA and 14 different inorganic salts and small organic molecules. These interactions were difficult to detect and visualize using gel electrophoresis with unbent DNA strands; however, our bent DNA bows were able to amplify these interactions, making them much easier to visualize and quantify. The amplification was facilitated by the bending energy in the bent DNA bows, which drove the conversion of the bent DNA bows to relaxed species, such as relaxed loops (i.e., straightened double-stranded segment) or dissociated single-strands ^41,76^. In addition, this technique based on bent DNA bows were capable of quantify the DNA interactions with the various inorganic salts and small organic molecules by fitting the relation between the amount of bent DNA bows and the concentrations of the tested salts or molecules presented in the solutions (i.e., *I*_B_ vs *c*) using the modified Hill equations (Eqs. 1 and 2). The strength of the interactions the tested salts and molecules with DNA can be reported by the characteristic concentration *u* in the modified Hill equation.

This work highlights the amplification effects of bent DNA bows on the interactions between DNA and other molecules. The demonstrations using 14 inorganic salts and organic molecules tested in this study suggested that the bent DNA bows could serve reliably as amplifiers and biosensors for many other DNA-interacting molecules. This work may also pave the way for exploiting the bent DNA bows for other applications such as monitoring water quality and screening DNA-targeting molecules and drugs ^77^. Furthermore, we expect that the bent DNA bows might be useful for understanding protein-DNA interactions by not only amplifying the interactions but also providing an additional controlling parameter (i.e., the curvature of DNA) ^78–81^.

The bent DNA bows focus on improving the sensitivity of DNA interactions with small molecules by amplification; however, the DNA bows are limited in distinguishing different ions or molecules. In other words, although the strengths of the interactions of different inorganic salts and organic molecules with the bent DNA bows are different, it is expected to be practically difficult to reliably map back to the type of ions/molecules from the strengths (i.e., *u* values or *I*_*B*_-*c* curves). Nonetheless, this limitation of our method may not be a concern in many situations such as screening DNA-targeting molecules or drugs in pharmaceutical applications ^82,83^.

Gel electrophoresis was used in this study for visualizing the interactions of the various tested salts and molecules with DNA. It is a commonly used, simple, economic biochemical technique, available in most biological, biochemical, and/or biophysical laboratories ^84–86^. One advantage of using gel electrophoresis to read out the signals of DNA interactions with other molecules amplified by the bent DNA bows lies in the accessibility, simplicity, economy and broad range ^85,86^, which is expected to make the bent DNA bows broadly useful. On the other hand, we should emphasize that the key role of our bent DNA bows is to amplify the interactions and thus generally improve the sensitivity of the original techniques; therefore, many other techniques are expected to be compatible with the bent DNA molecules for measuring and visualizing the signals of DNA interactions with other molecules. It would be interesting to extend the visualization methods to others, such as techniques based on fluorescence, melting temperature, calorimetry, circular dichroism, Raman spectroscopy, and nuclear magnetic resonance, and investigate how the bent DNA bows enhance the sensitivity of these techniques.

## Acknowledgments

This work was supported by the University of Arkansas, the Arkansas Biosciences Institute (Grant No. ABI-0189, No. ABI-0226, No. ABI-0277, No. ABI-0326), the Arkansas Department of Higher Education (Grant No. 003049-00001A), the Center for Functional Nanomaterials at Brookhaven National Lab (Grant No. 38596), and the National Science Foundation (Grant No. 1826642).

## Author contributions

YW conceived the project; JF and MR carried out the gel electrophoresis experiments; LZ and SW carried out the TEM experiments; JF, LZ and YW performed the data analysis; JF, LZ and YW wrote the manuscript. All authors reviewed and revised the manuscript.

## Supporting Information

**Figure S1.**
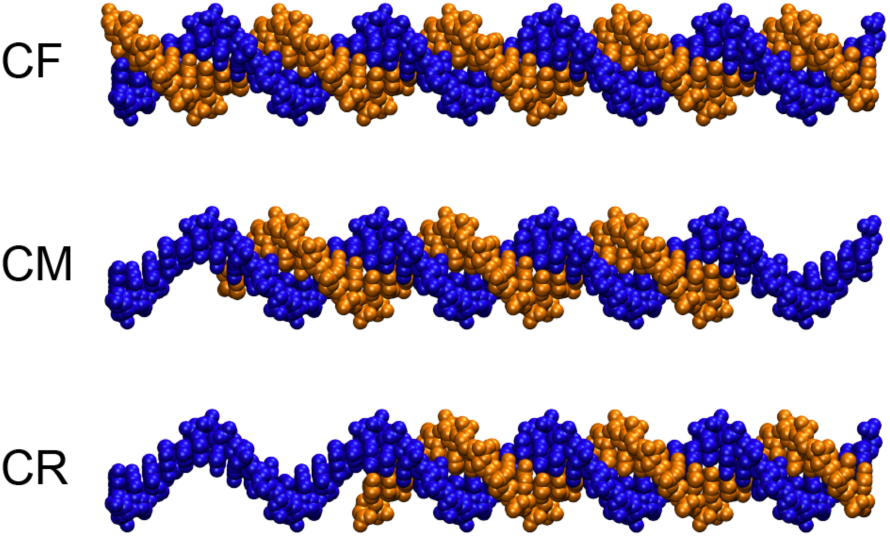
Linear DNA as negative controls: (A) fully double-stranded DNA, (B) partially double-stranded DNA in the middle, and (C) partially double-stranded DNA in the right.

**Figure S2.**
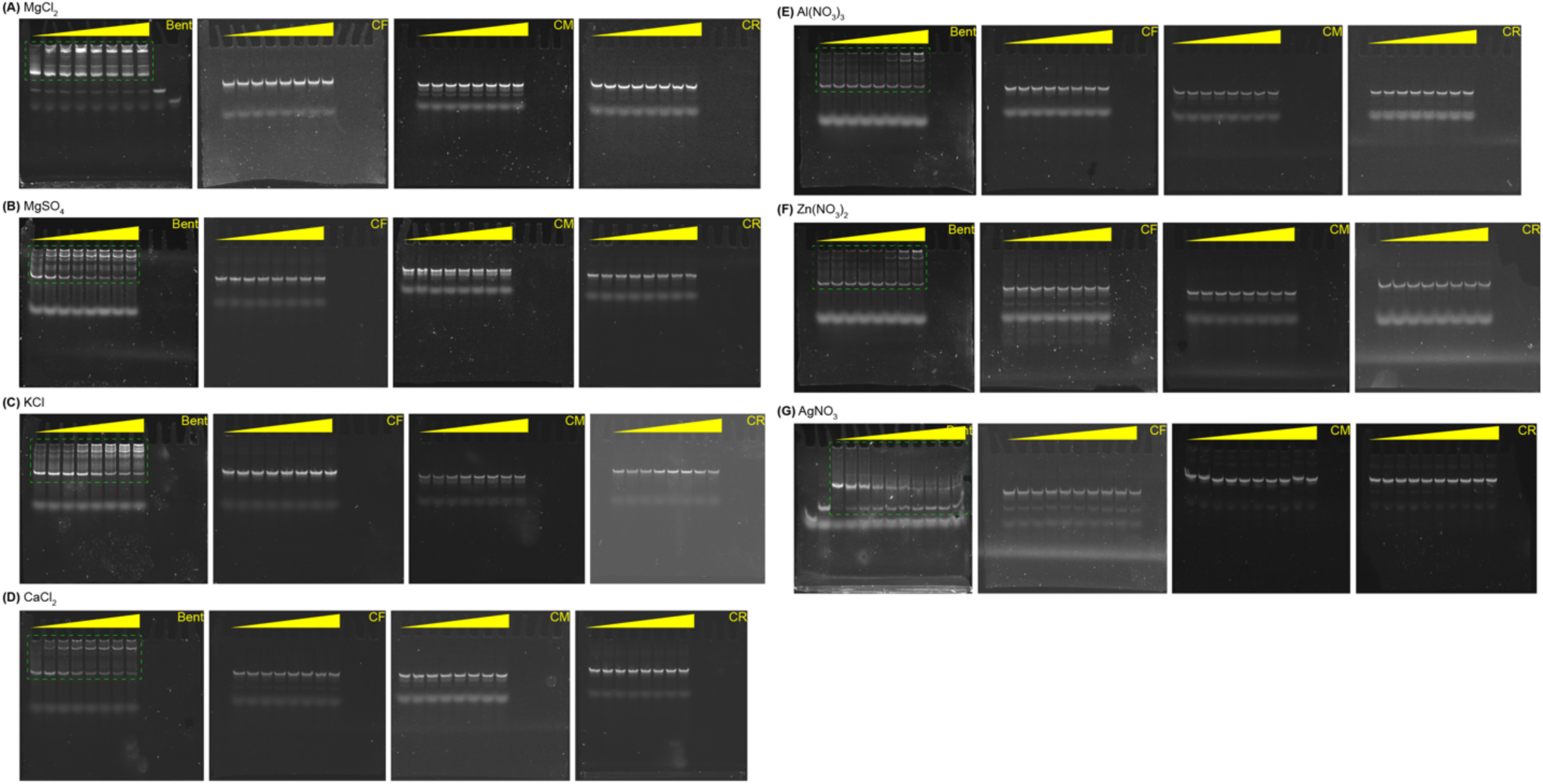
Examples of full-length gels for DNA bows or linear DNA in the presence of salts at increasing concentrations: (A) MgCl_2_, (B) MgSO_4_, (C) KCl, (D) CaCl_2_, (E) Al(NO_3_)_3_, (F) Zn(NO_3_)_2_, and (G) AgNO_3_. Green rectangles indicate the cropping areas of the gels for the insets in Fig. 3.

**Figure S3.**
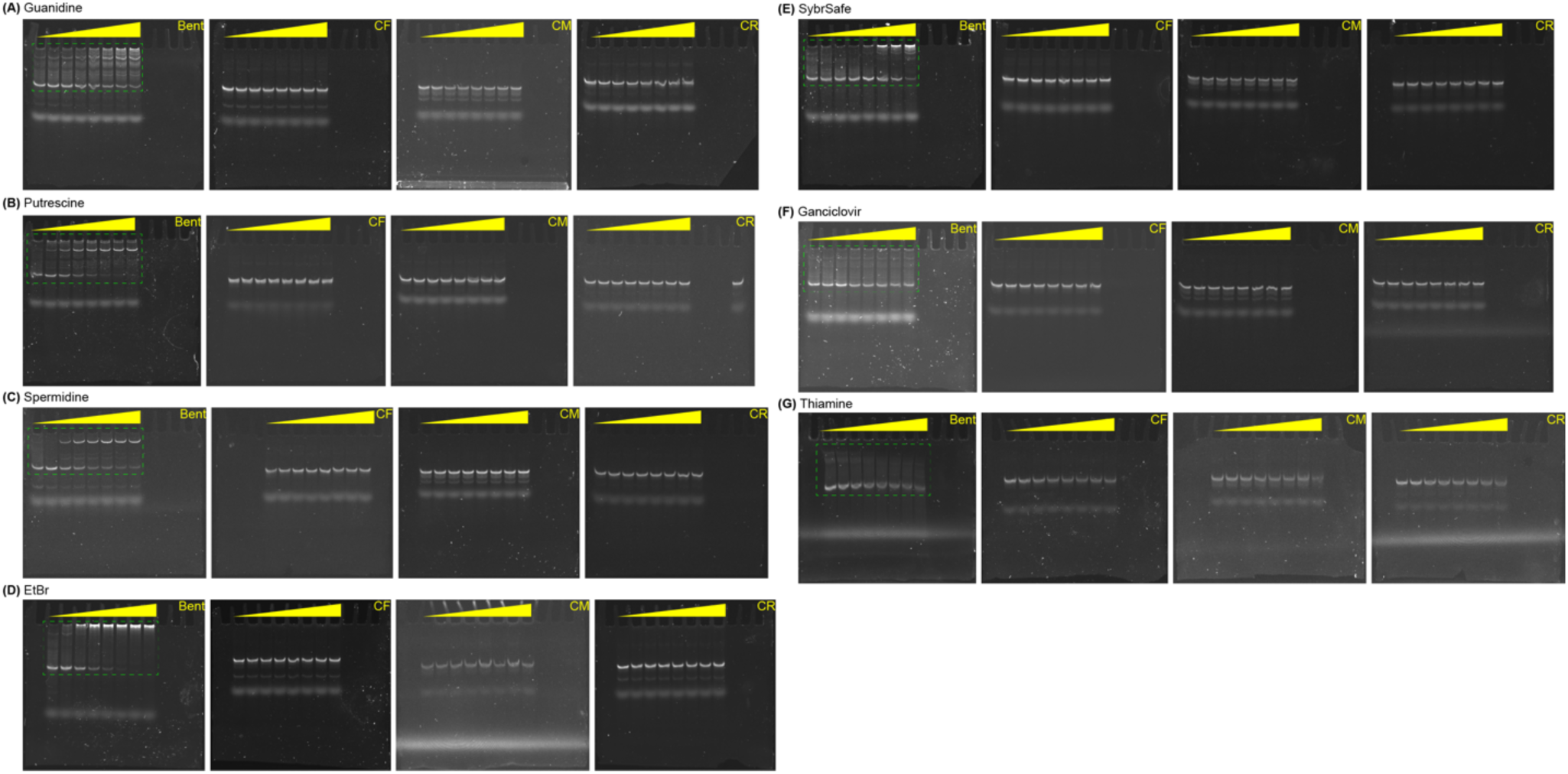
Examples of full-length gels for DNA bows or linear DNA in the presence of small organic molecules at increasing concentrations: (A) guanidine, (B) putrescine, (C) spermidine, (D) ethidium bromide, (E) SYBR safe, (F) ganciclovir, and (G) thiamine. Green rectangles indicate the cropping areas of the gels for the insets in Fig. 4.

**Figure S4.**
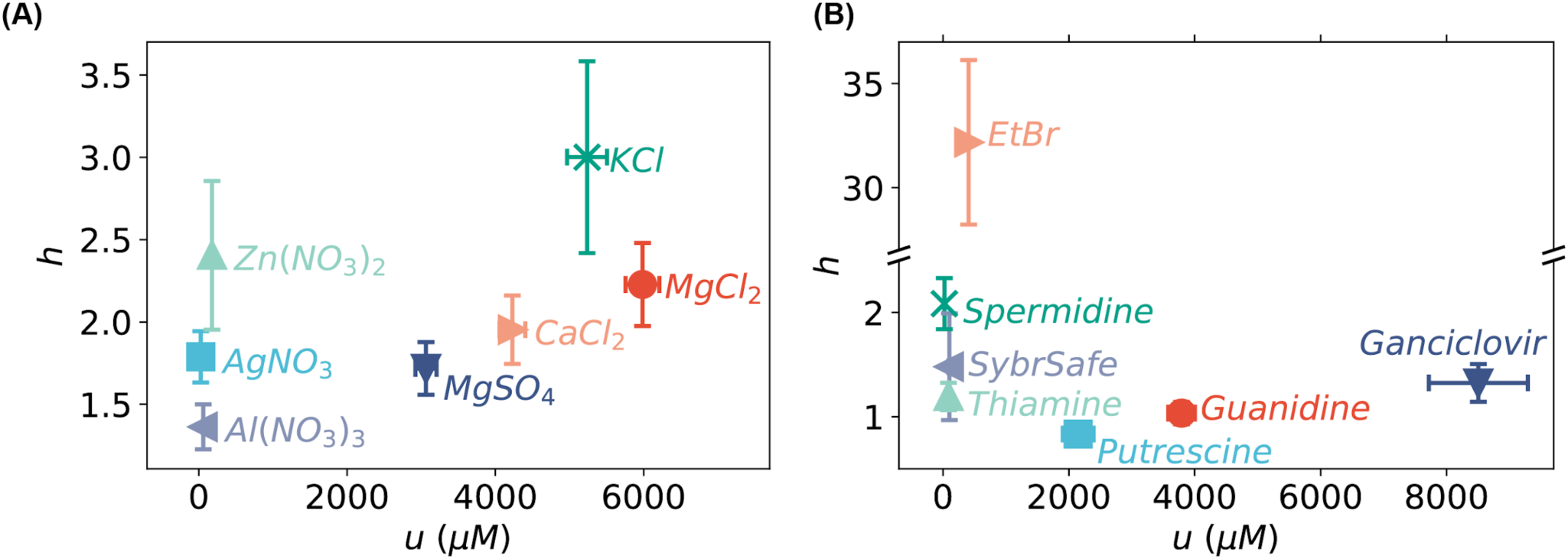
Fitted *h* and *u* values from the modified Hill equation (Eq. 1) for the (A) inorganic salts and (B) organic molecules.

**Figure S5.**
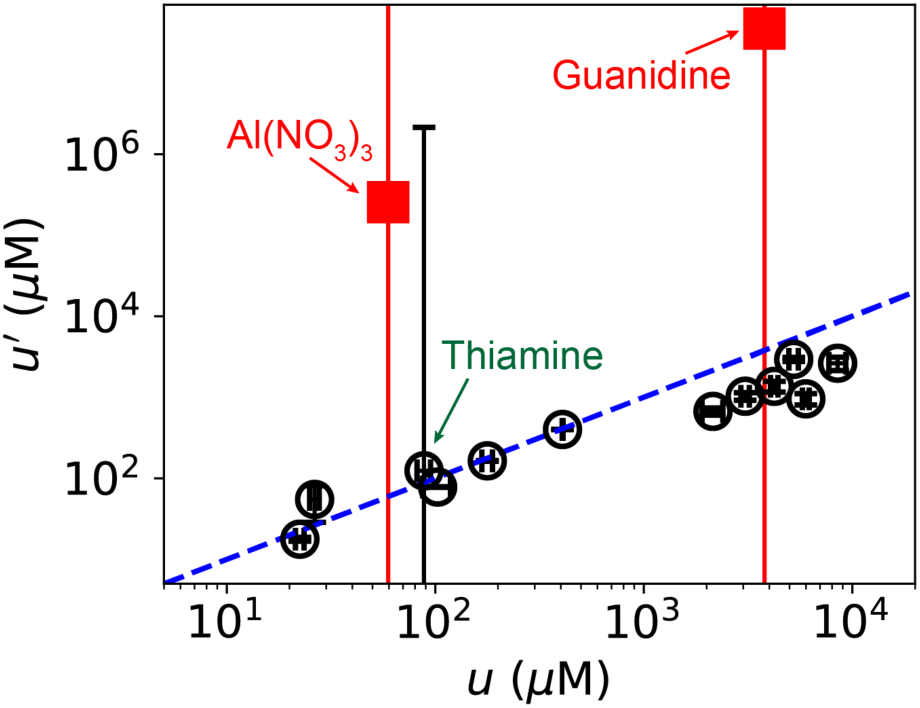
Correlation between *u*′ (fitted from relaxed DNA) and *u* (fitted from bent DNA bows). Error bars represent fitting errors. The blue dashed line indicates *u*′ = *u*.

